# Insights into the triggering factors and determination mechanisms of host alternation in aphids

**DOI:** 10.1101/2024.09.20.613903

**Authors:** Aurelie Etier, Rituparna Ghosh, Théo Vericel, Romuald Cloteau, Gaëtan Denis, Maïwenn Le Floch, Yannick Outreman, Gaël Le Trionnaire, Jean-Christophe Simon

**Affiliations:** UMR 1349 IGEPP, INRAE, Institut Agro, Université Rennes 1, Le Rheu, France; UMR 1349 IGEPP, INRAE, Institut Agro, Université Rennes 1, Rennes, France

**Keywords:** *Rhopalosiphum padi*, *Prunus padus*, Nutritional polyphenism, Primary and secondary host, Morph induction

## Abstract

The bird cherry-oat aphid *Rhopalosiphum padi*, is one of the ten percent of aphids that are host alternating. The annual life cycle of *R. padi* entails an obligatory migration between a primary woody host, *Prunus padus* (Rosaceae), where sexual reproduction occurs, and many secondary hosts of the Poaceae family, where successive generations reproduce parthenogenetically. Migration between the two types of host plants is achieved by specialized migrant morphs. Very little is known about the cues that trigger migrant development, host alternation establishment and the underlying mechanisms. In this study, we aimed to elucidate the key steps involved in migrant induction and determination of host alternation, and to shed light on these intricate processes. We showed that a specific stage of leaf maturity triggers the development into a migrant morph when presented to *R. padi* at the first or second larval instar stages. We also demonstrated the gradual establishment of acclimation to the secondary hosts that precedes the migration step. Finally, we showed that the switch to feeding on the alternative hosts is irreversible once emigrants have reached adulthood. These results not only allowed us to define the critical periods of the leaf stage associated with morph induction and determination, but also to pinpoint the emigrants’ developmental stages when acclimation to secondary hosts begins. These novel insights of these critical periods will facilitate the investigation of molecular changes associated with host alternation in aphids.

## 1. Introduction

Aphids, belonging to the hemipteran order and the family Aphididae, represent a diverse group of insects comprising more than 4700 identified species (Blackman & Eastop, 2007). The phenotypic plasticity exhibited by aphids enables them to quickly cope with varying seasonal, crowding, or dietary conditions, thereby facilitating their remarkable capacity to colonize and utilize plants (Blackman, 2000). A striking illustration of this plasticity is their ability to produce alternative phenotypes in response to fluctuating environmental factors, a phenomenon referred to as polyphenism (Moran, 1992). Environmental cues serve as pivotal triggers for polyphenism, initiating phenotypic divergence through neurochemical and hormonal pathways that induce short-to long-term transcriptional reprogramming, often mediated by epigenetic regulatory mechanisms (Srinivasan & Brisson, 2012). Polyphenism in aphids can be categorized into four main types: caste (defensive vs reproductive morphs), reproductive (sexual vs clonal morphs), dispersal (winged vs wingless morphs), and nutritional (Braendle et *al*., 2006; Shibao et *al*., 2010; Yan et *al*., 2020). The latter enables them to transition typically from woody host plants (during winter) to herbaceous host plants (during summer), a phenomenon also known as host-alternation (Blackman & Eastop, 2007; Dixon, 1998). It has been proposed that host-alternation represents an adaptation to seasonality by allowing nutritional complementation between hosts whose quality and presence/availability fluctuate across the seasons (Dixon, 1998). However, the fact that only ten percent of all aphid species can alternate hosts has prompted certain authors to suggest that rather than being adaptive, host alternation may be a consequence of evolutionary constraints (Moran, 1992; Blackman, 2006; Blackman, 2000). Aphids pose a significant threat to crop yield directly by taking up plant nutrients or indirectly by transmitting phytopathogenic viruses. Although many economically important aphids are host alternating species (e.g. *Aphis fabae, Myzus persicae, Rhopalosiphum padi*), the cellular and molecular mechanisms underlying host alternation in aphids are currently unknown. Therefore, gaining a deeper comprehension of host alternation is imperative for mitigating the economic impacts of aphid infestations on crops.

Among host-alternating species, *Rhopalosiphum padi* stands out as a prominent cereal pest in Europe, primarily attributed to its role in the transmission of Yellow Dwarf Viruses of Cereals (Jarošová, Żelazny, & Kundu, 2019). *Rhopalosiphum padi* undergoes host alternation, transitioning between a primary woody host within the *Prunus* genus during winter (*Prunus padus*, exclusively in Europe) and numerous secondary hosts belonging to the Poaceae family during summer (e.g. wheat, barley, oat, maize, ray-grass and numerous wild grasses) (Dixon, 1971b). The life cycle of *R. padi* begins with oviposition on *P. padus* in the early winter (see Supplementary data figure S1). Wingless fundatrix emerges from overwintering eggs in the early spring. The fundatrix produces wingless *fundatrigeniae* by parthenogenesis (asexual reproduction). Both fundatrix and *fundatrigeniae* are specialized to the primary host. After two to three parthenogenic generations of *fundatrigeniae*, spring emigrants appear, which alternate hosts (Wiktelius, 2009). Emigrants colonise the secondary hosts and produce wingless *virginiparae*, which are specialized to the secondary hosts. During the summer, both wingless and winged *virginiparae* are produced on the secondary hosts. In response to the shortening of day length, two different winged morphs, the *gynoparae* and the male, are produced that will make the reverse migration from the secondary to the primary host. On the primary host, sexual reproduction occurs between male and the *oviparae* (wingless sexual female produced by the *gynoparae*), to produce eggs.

Upon the spring migration, environmental cues are perceived by the *fundatrigeniae* that induce emigrant development (see Supplementary data figure S1). Aphid development consists of four larval stages followed by an adult stage. The first and second larval stages (L1 and L2) of *fundatrigeniae* and emigrant are morphologically identical. However, L3 and L4 (*fundatrigeniae*) differ morphologically from N3 and N4 (emigrant) stages, with visible wing buds in the latter. In the *fundatrigeniae*, L3 develops into a wingless adult that remains on *P. padus*. In contrast, in the emigrant, N3 develops into a winged adult that feeds on *P. padus* until adulthood before being repelled by the primary host and moves to the secondary hosts (Dixon, 1971b; Glinwood & Pettersson, 2000).

The nutritional polyphenism in *R. padi* is influenced by changes in environmental conditions, such as crowding and plant-emitted signals (Dixon & Glen, 1971; Wiktelius, 2009). Dixon and Glenn (1971) underscored the significance of plant signals in triggering nutritional polyphenism in *R. padi*, specifically by showing the involvement of plant maturity. However, this study does not provide clarity regarding the specific age or maturity level of the leaf responsible for instigating this morph switching. To address this knowledge gap, a time-course analysis is essential for precisely delineating leaf age/maturity of the leaf responsible for triggering the development of the emigrant.

We also lack information regarding the temporal window during larval development wherein perception of plant-emitted signals and/or crowding trigger the development of emigrants. It remains unclear whether these signals can be detected and responded to as early as the first generation of fundatrigeniae, or if a few generations must pass before the development of emigrants. Besides, as previously mentioned, emigrant development becomes apparent from the third larval instar stage, indicating two potential sensitive time frames. The first possibility involves a pre-natal sensitive period when aphid embryos are still within the mother’s abdomen, facilitating the transmission of signals from the mother to her offspring (Dixon, 1998). The second possibility corresponds to a post-natal sensitive period, where signal perception occurs during the first and/or second larval instar stages and initiates the switch in only a few days.

Moreover, our understanding of the establishment mechanism of nutritional polyphenism during emigrant development remains elusive. Previous investigations have primarily focused on this process in emigrant adults, revealing a gradual shift in host preference, notably observed as a heightened preference for secondary hosts in 24 to 48 hours aged emigrant adults compared to newly emerged individuals (Glinwood & Pettersson, 2000). In aphids, there is a well-established phenomenon of mutual exclusivity between flight development and settling on the secondary hosts (Dixon, 1971a). This suggests a sequential activation pattern, wherein flight development precedes acclimation to the new host. However, it is apparent that aphids can only encounter the new host once they possess the ability to fly and locate it. This does not imply that the process of host acclimation initiates solely after the aphid reaches the secondary host; rather, such a delay would compromise the aphid’s fitness on the secondary host. Consequently, it can be postulated that the developmental machinery governing host switching and acclimation operates in parallel rather than in series. Further investigation is warranted to elucidate the initiation of nutritional polyphenism during larval development. It is conceivable that a complex dynamic interplay of molecular mechanisms involving flight readiness and host acclimation exists, raising the possibility of reversing host switching. Host switching may be imperative in natural habitats to counteract the adverse effects of summer *P. padus* leaves; however, in the persistent presence of spring leaves, aphids may remain on the primary host. The possibility of trapping *R. padi* on the primary host will be of particular interest for agricultural benefit.

With this background, the objectives of this study were to precisely define the key steps of the nutritional polyphenism in *R. padi*. To address this, we formulated four questions: (i) What is the critical period of leaf maturity which is sufficient to induce emigrant production switching from the primary host to the secondary host? (ii) At which generation / larval stage the *R. padi* larvae can sense the signals, enabling the development of emigrants? (iii) Is there a gradual establishment of nutritional polyphenism in *R. padi*? (iv) Could host switching in young adults be a reversible mechanism in response to perceiving favorable conditions on primary hosts?

## 2. Materials and Methods

### Insect and plant sources

Ten *Rhopalosiphum padi* fundatrices at the L1/L2 developmental stages (i.e. few days after egg hatching) were collected on *Prunus padus* in the beginning of March 2023 from Le Rheu city, France (48.1011° N, 1.7954° W). Fundatrices were individually reared immediately upon collection onto young leaves in isolation within clip-cages (1.5cm diameter) to mitigate potential crowding effects, which could interfere with the perception of plant cues. They were placed onto the back of uninfested leaves. Young *P. padus* plants of 75-100 cm hight were purchased from local nurseries. These saplings were planted singly into pots (3 litres) with soil (60% white peat, 40% sand, 14% coconut fibre medium, 6% clay). Wheat (T*riticum aestivum*, geny variety) seeds were used as the secondary host and were germinated individually in the same soil inside 15 mL plastic tubes with holes for water penetration. 15-days old seedlings were used for experiments. Plants and insects were grown inside climatic chamber with constant temperature (18°C) and photoperiod [16:8 h (L:D)], to eliminate seasonal effect. The plants were used for growth measurements, insect culture maintenance and assays. No fertilizers or pesticides were added after planting to avoid any change in plant growth.

### Defining leaf age

*Rhopalosiphum padi* emigrants are produced in late spring. It was hypothesized that change in leaves’ biochemistry is responsible for this winged morph induction. To mitigate the environmental impact on this phenomenon, plants were grown at constant temperature and photoperiod. Leaf growth was monitored by precisely measuring their length (mean ± SE; in mm). Only the first flush of leaves was used in this experiment to avoid high variance in data due to (1) nutrient deficiency with time, and (2) influence of already existing leaves on the growth of second or next flushes of leaves. Measurements of the same leaves (n= 8) were noted, three times a week, over a period of 30 days post bud-burst.

### Wing induction experiment

From five fundatrices reared in clip-cages, the first four individuals produced by each fundatrix were collected at the first larval instar (L1). The larvae were introduced to two contrasted categories of leaves based on the growth pattern analysis of *P. padus* leaves: 15 days-old (considering the bud-burst event as day zero) leaves (hereafter referred to as young leaves) and 30 days-old leaves (hereafter referred to as mature leaves) (see Supplementary data figure S2 for a graphical representation of the experimental protocol). The larvae were distributed as follows to the two conditions: out of the four larvae colllected from one fundatrix, two were deposited within clip cages onto young leaves (referred to as condition 1) and two onto mature leaves (referred to as condition 2). Larvae reared exclusively on young leaves until reaching the L3 stage were maintained in clip cages on young leaves until larviposition. From one fundatrigeniae, two L1 were then subjected to two conditions: one in Condition 3 placed on mature leaves immediately after larviposition, and one in Condition 4 initially maintained on young leaves and transferred to mature leaves after reaching the L2 stage. In total 20 freshly born larvae were deposited individually for each condition as replicates.

After reaching the third larval instar (4 days after transfer) the ratio of L3/N3 for all the tested conditions, which represents the proportion of future *fundatrigeniae* vs emigrants, was monitored under binocular (magnification 6x).

### Choice and no-choice preference tests

From the fundatrices maintained in clip-cages, 100 fundatrigeniae were reared on actively growing plant for one generation. The second generation was exposed to mature leaves, and were collected at six stages: (i) L1, (ii) L2, (iii) N3, (iv) N4, (v) winged adults before leaving *P. padus* leaves, and (vi) emigrant 24h after leaving *P. padus* leaves. A settling choice bioassay was used to determine host plant preference at each stage. Whole-plants for *P. padus* and 15 days old wheat seedlings were used. One mature leaf of *P. padus* still attached and two-branch tip of wheat leaves were placed on each side of a Petri dish (⊘ 90mm). The Petri dish was covered with a lid and sealed with parafilm. Additionally, the wingless fundatrigeniae specialized to primary host and wingless and winged virginiparae specialized to secondary hosts were used as controls to validate our experimental approach (i.e. we expect wingless fundatrigeniae to be more attracted by *P. padus* than wheat and the opposite for virginiparae). To stimulate movements, aphids were starved for two hours, then a total of 20 individuals per stage were released into the centre of the Petri dish and the experiment was repeated six times. The plant on which they settled was recorded after 12 hours, a time which is generally enough for choice establishment in aphids (Sochard, Dupont, Simon, & Outreman, 2021; Sochard, Le Floch, Anton, Outreman, & Simon, 2021).

### Survival study of the emigrants on wheat

From the 100 fundatrigeniae reared on actively growing plant for one generation, the produced larvae were collected at six stages after exposition to conditions inducing emigrant production (mature leaves): (i) L1, (ii) L2, (iii) N3, (iv) N4, (v) winged adults before leaving *P. padus* leaves, and (vi) emigrant 24h after leaving *P. padus* leaves. Forty individuals per stage were individually transferred and reared on one single 15-days old wheat seedling. After wheat infestation, survival was monitored daily for seven days.

### Testing the reversibility of the host alternation in *R. padi* emigrant adults

Hundred fundatrigeniae were reared on actively growing *P. padus* plant for one generation. Their progeny was reared onto mature leaves and were collected as winged adults before leaving *P. padus* leaves stage and individually placed into plastic bags containing one leaf either young or mature or one single 15-days old seedling. A total of 40 adults were placed per bag. Survival rates of the *R. padi* adults were monitored daily for a period of seven days to investigate the viability and the reversibility of the host alternation.

### Data analysis and statistics

All the statistical analyses were performed using R software R v4.2.2, with value of p < 0.05 considered as significant.

The statistical tests and analyses employed in this study are detailed in the supplementary data section.

Results of GLM, GLMM and Cox hazard survival model are summarized in supplemental data table S1, S2, and S3.

## 3. Results

### 3.1. Characterization of the leaf growth pattern of *P. padus*

*Prunus padus* leaves followed a growth pattern, which could be divided into four phases based on their length (Figure 1): (1) initial slow growth phase with a growth rate of 1.9mm/day up to fourth day (7.75 ± 0.96mm), (2) fast growth phase with an average growth rate of 5.7mm/day from fifth to 19^th^ day (94.5 ± 4.73mm), (3) final slow growth phase with an average growth rate of 2.1mm/day from 19^th^ to 23^rd^ day (103.25 ± 2.88mm), and (4) very slow/ no growth phase. With these results, 15-days and 30-days old leaves were selected as representatives of young and matured leaves, respectively.

**Figure 1.**
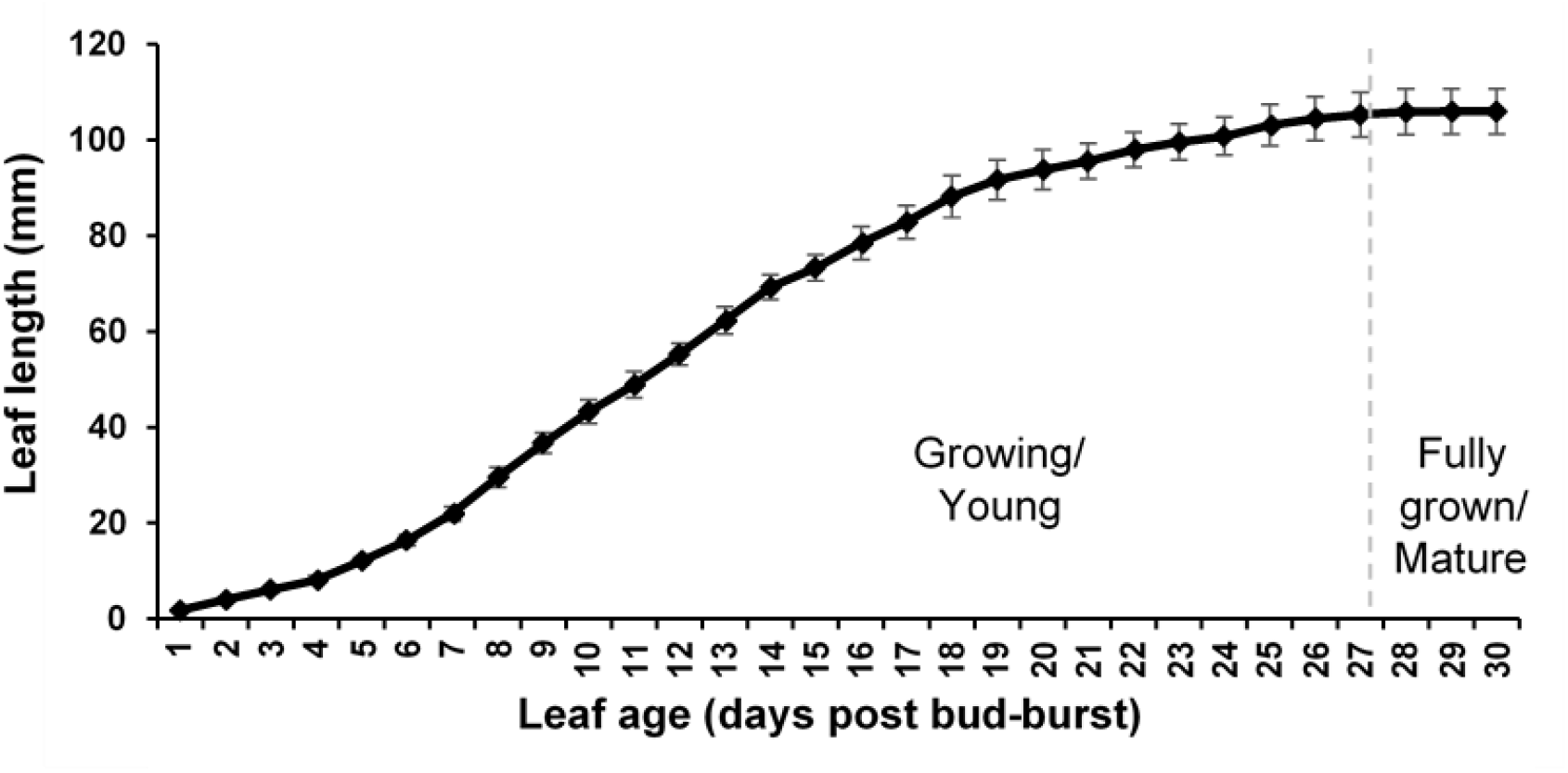
*Prunus padus* leaf growth kinetics. The line (mean ± SE) represents the leaf length (n= 8 leaves from three plants). Growing and mature leaves were used for further experiments.

### 3.2. The role of leaf maturity in inducing emigrant development and the responsiveness of generations and first and second instar larvae

Based on the growth pattern analysis, we hypothesized that young leaves would induce the production of fundatrigeniae, while mature leaves would induce emigrants. Freshly born first instar larvae placed onto young leaves (condition 1) developed into 100% fundatrigeniae. Conversely, the majority (90%) of L1 reared on mature leaves (condition 2) developed into emigrants (Figure 2). The effect of leaf age on winged/wingless proportion was highly significant (χ^2^ = 12.8, df = 1, p < 0.01) and the maturity of leaves was sufficient in inducing emigrants’ development. The L1 from the first (condition 2) or second (condition 3) generation following fundatrix stage showed no significant difference in their responsiveness to plant maturity cues (binomial GLMM, χ^2^ = 63,823, df = 2, p=0.88). Additionally, larvae exposed to mature leaves during the first instar (condition 3) or during the first and second instars (condition 4) did not reveal a significant difference in the morph determination (binomial GLMM, χ^2^ = 63,823, df = 2, p=0.31). These findings indicate that both first and second generations after fundatrix are sensitive to leaf maturity exposure. Furthermore, in the second generation, L1 and L2 were both able to perceive the leaf-emitted signal, leading to the development of emigrants (Figure 2).

**Figure 2.**
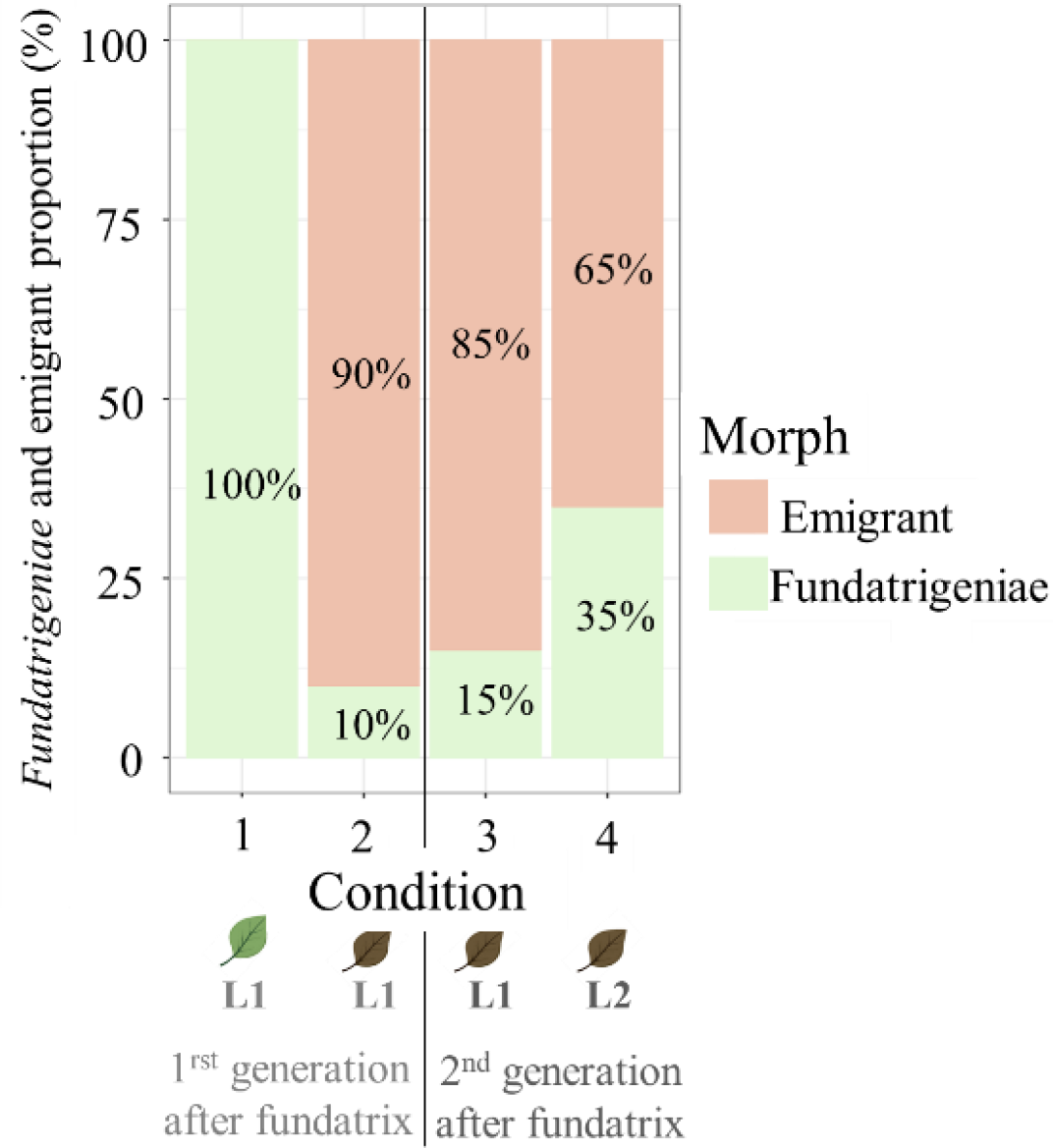
Influence of the plant maturity on wing induction. Monitoring of wing induction under four experimental conditions. Condition 1 involved placing individual L1 larvae produced by fundatrix in clip cages on young leaves, while Condition 2 entailed placing them on mature leaves. Furthermore, L1 larvae from the second generation (after Condition 1) were subjected to two additional conditions: transfer to mature leaves immediately after birth (Condition 3) or after reaching the L2 stage (Condition 4).

### 3.4. Nutritional polyphenism is gradually established during the larval development in *R. padi*

#### 3.4.1. The initiation of the switch in the preference of host-plant

In order to investigate the temporal dynamics of nutritional polyphenism concurrent with emigrant development, the preference for primary and secondary hosts was evaluated. We found in our host plant choice assay that the mean proportion of *fundatrigeniae* opting for wheat was approximately 5%, whereas more than 85% of secondary host-specialized wingless/winged adults exhibited this preference, consistent with anticipated outcomes and showing the relevance of our experimental device. Due to a high mortality rate observed after the manipulation and starvation of the first larval instar upon their deposition in Petri dishes (98%, data not shown), the analysis of their preferences became unfeasible. Furthermore, the different stages or morphs significantly affected the preference for wheat in *R. padi* (binomial GLMM, χ^2^ = 132.32, df= 7, p < 2.2e-16) (Figure 3), with *fundatrigeniae* again serving as the reference. The GLMM’s coefficients (Supplementary data table S1) showed that *fundatrigeniae* and the emigrants at the second and third larval instars had a similarly very low preference for wheat. However, the wheat preference in fundatrigeniae was significantly lower than for the fourth larval instar onward, but no significant difference with the third larval instar. These results indicate an initiation of host plant preference switch from the fourth larval stage.

**Figure 3.**
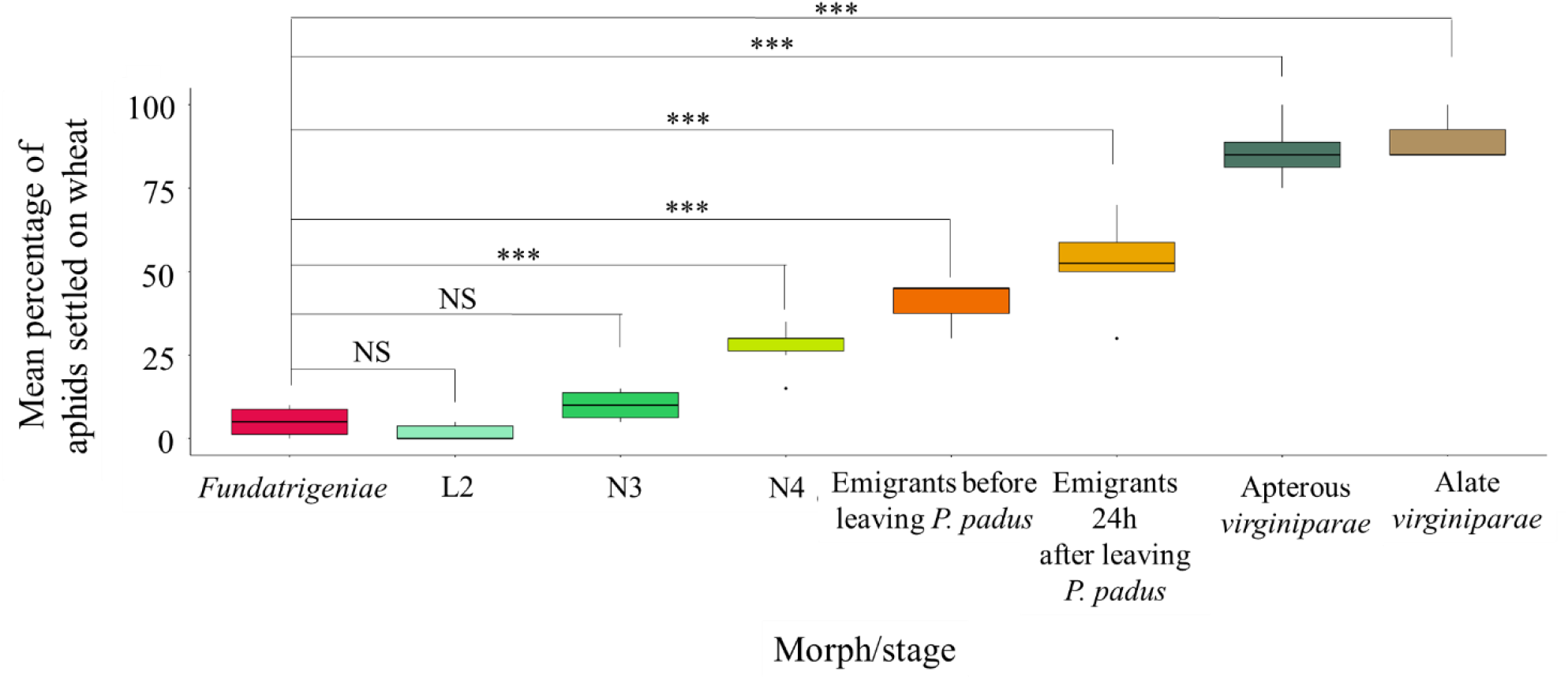
Proportion of *Rhopalosiphum padi* individuals at various morph or emigrant developmental stages selecting wheat as the host plant when presented with a choice between *P. padus* (primary host) and wheat (secondary host). Significant differences compared to the *fundatrigeniae* are labeled as NS for “non-significant”, *p < 0.05, **p < 0.01, and ***p < 0.005.

#### 3.4.2. The initiation of the pre-acclimation to wheat

The last observations support the hypothesis of a gradual establishment of host switching, potentially occurring from the fourth larval instar onwards. To investigate possible pre-acclimation to wheat feeding, the settling behavior of emigrants was studied by monitoring their survival rate daily for seven days after their transfer to wheat at the six stages described previously. The different stages significantly affected the survival of *R. padi* on wheat (Cox proportional hazards model, χ^2^ = 68.46, df = 5, p < 0.001) (Figure 4). Of all the different stages tested, the first and second larval instars exhibited significantly lower survival rate, with approximately half of the individuals that were dead 24 hours after transfer to wheat (Supplementary data table S2). The stage N3 displayed a moderate survival rate upon the seven days, significantly higher than that of the L1 and L2 stages. The survival rate for the N4 stage did significantly differ from that of the N3 stage and was comparable to that of the emigrants before leaving *P. padus* and the emigrant after the flight. These findings indicate a gradual establishment of secondary host pre-acclimation initiated from the third larval instar and firmly established from the fourth one.

**Figure 4.**
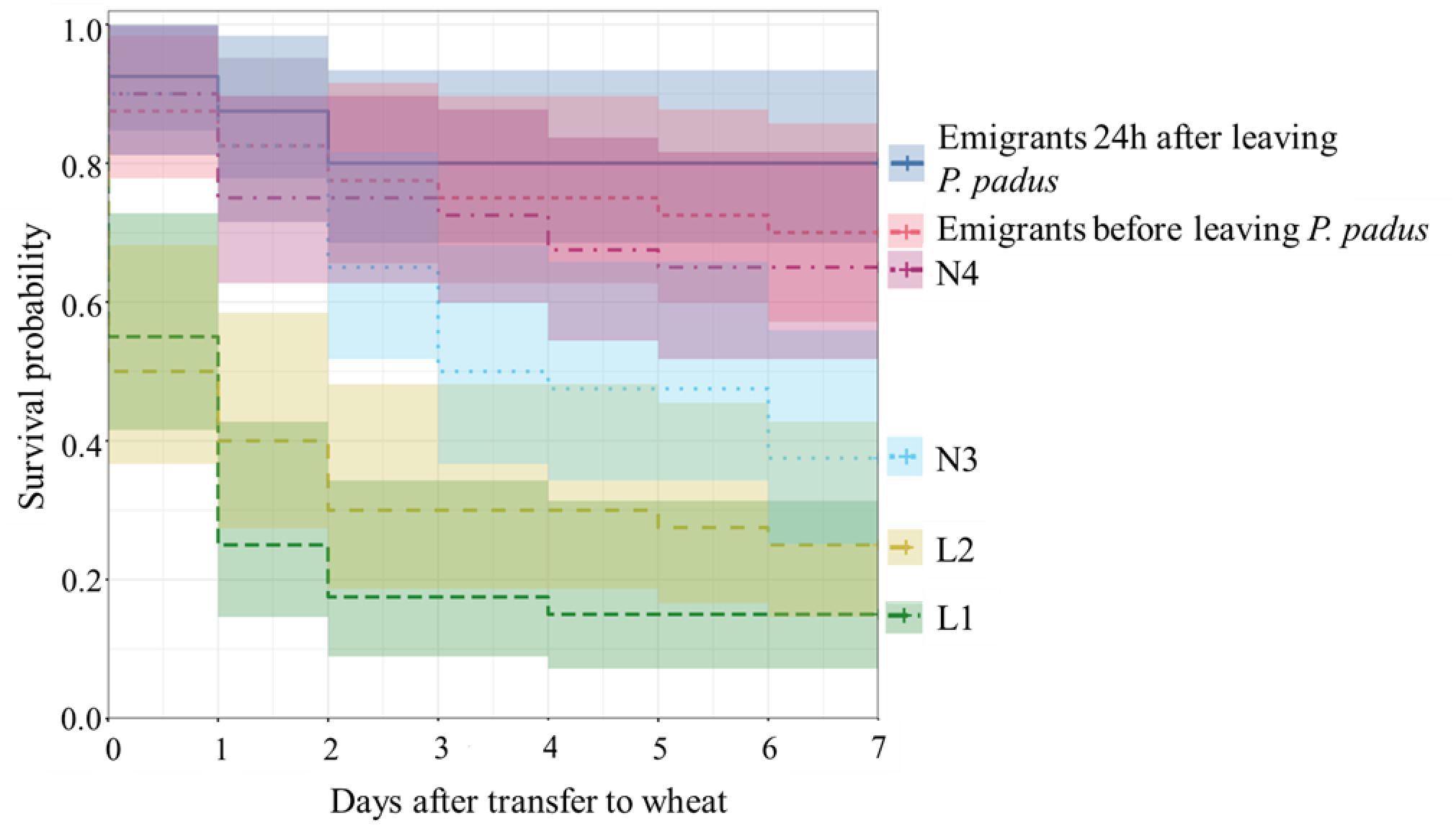
Estimated survival probability of *R. padi* individuals installed on wheat at six different emigrant developmental stages. Each vertical step in the curve indicates a death event.

### 3.6. Host switching is irreversible in adult emigrants

The potential reversibility of host alternation following exposure to new favorable conditions of *P. padus* was investigated in this experiment. Within four days after transfer, half of the emigrants installed on either young or mature leaves of *P. padus* were dead. Furthermore, a significant reduction in survival rate was observed in emigrant adults reared on either young or mature leaves compared to those reared on wheat (80%) (Cox proportional hazards model, χ^2^= 27.86, df = 2, p < 0.001) (Figure 5). In addition, there was no significant change in the survival rates between individuals reared on young or mature leaves of *P. padus* (Supplementary data table S3). These findings underscore the irreversibility of host switch, even under conditions favorable for the growth of *R. padi* adult emigrants, such as young leaves of *P. padus*.

**Figure 5.**
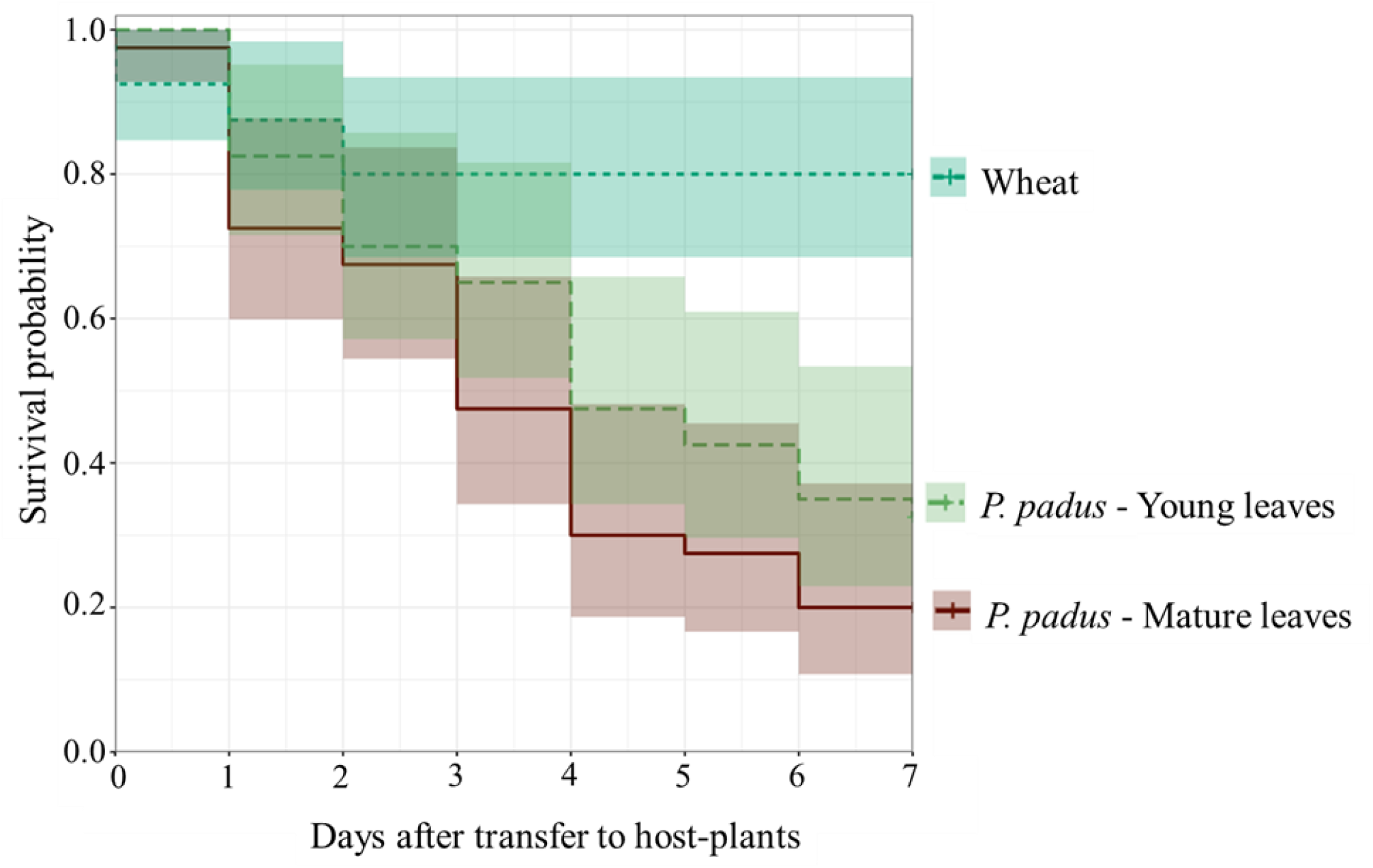
Estimated survival probability for emigrants of *R. padi* (collected before leaving *P. padus*) reared on either young leaves (light green) or mature leaves of *P. padus* (brown) or on wheat (dark green). Each vertical step in the curve indicates a death event.

## 4. Discussion

In this study we first characterized the growth pattern of *P. padus* leaves, providing a proxy for assessing leaf maturity. We subsequently demonstrated that the leaf maturity signal, perceived in the first and second larval instar, is sufficient to trigger the development of emigrants. Furthermore, we showed that the onset of feeding change is a gradual and irreversible process that appears to be initiated at the fourth instar upon the development of the emigrants.

### Leaf maturity alone triggers migrant development

In our study, two distinct leaf growth stages were identified: 15-day-old young leaves and 30-day-old mature leaves. As leaves mature, they transition from being metabolite sinks to sources, which affects their nutritional quality (Coleman, 1986). Insects’ preference-performance highly depends on plant chemistry, which undergoes changes upon leaf maturation and associated biochemical alterations, influencing the plant-insect relationship (Barber & Marquis, 2011; Feeny, 1970). This change includes a reduction in essential nutrients and possibly an increase in toxic compounds, making the leaves less suitable for insect consumption (Coley, 1980; Haukioja, Ossipov, & Lempa, 2002). Consequently, aphids might respond to the decline in food quality by developing into winged emigrants to seek better habitats. Our findings showed that this transition from young to mature leaves alone was sufficient to trigger this emigrant development. This suggests that aphids can rapidly detect changes in leaf maturity and adjust their development accordingly, highlighting the importance of leaf quality in aphid colony survival and dispersal. In case of *P. padus*, the leaf maturation coincides with changes in amino acids and phenolics (Czerniewicz et *al*., 2011; Sandström & Pettersson, 2000), while the alteration of other compounds remains unexplored. Further research is needed to identify the specific plant compounds responsible for inducing this response.

### First and second larval instars are the critical stages to induce the migrant

The morph-switching mechanism relies on a sensitive period of larval development, identified here as the first and the second larval instar. Upon the nutritional polyphenism establishment, the production of wings is the first noticeable change. Wing development in aphids is considered the default developmental pathway, with wing primordia present in the first larval instar. During post-embryonic development, these wing primordia will either degenerate or continue their development, depending on environmental and physiological cues. The degeneration of wing primordia typically occurs during the second instar larvae stage (Ishikawa and Miura 2009). This explains that the window of sensitivity mainly resides at the first or second larval instars when exposed to mature leaves. In addition, our results suggest that emigrant determination occurs post-natally, as the mothers and their offspring were reared in clip-cages to avoid crowding effects. Although a maternal effect may also initiate this process, it is not mandatory in *R. padi* (Dixon and Glen 1971; Müller, Williams and Hardie 2001). Taken together, these findings highlight the rapid induction of nutritional polyphenism enabling the exploitation of two complementary seasonal food resources (Dixon, 1971a). This ability allows *R. padi* populations to anticipate deteriorating conditions, thereby ensuring the sustained survival of the colony.

### Acclimation to the secondary host is established during the larval development and is irreversible

In a prior study, it was demonstrated that the emigrant stays on *P. padus* from larval emergence until early adulthood, after which it is repelled from the primary host, requiring time for flight, before being attracted to secondary hosts (Glinwood and Pettersson 2000). It was hypothesized that emigrants should be fully developed before initiating settlement on secondary hosts (Leather and Dixon 1982). However, our study demonstrates an early preference and pre-acclimation of *R. padi* to the secondary host. Indeed, host choice tests showed gradual changes in preferences throughout emigrants’ development from the fourth larval stage. Existing literature suggests that the interplay between repulsion and attraction, guided by chemical cues, influences host plant preference (Anton & Cortesero, 2022). Volatiles such as methyl salicylate from *P. padus* play a key role in primary host repellence (Glinwood & Pettersson, 2000; Pettersson et al., 1994). The shift in preference may be due to repellent compounds produced by maturing *P. padus* leaves. The duration of flight before settling on wheat might be influenced by chemosensory pathway activation or repression for different hosts during emigrant development (Peng et al., 2021). Recent studies show that aphid host-plant adaptation involves molecular mechanisms that modulate plant defenses or detoxify host-specific secondary metabolites (Shih, Sugio, & Simon, 2023). The current study further reveals a predetermined capacity for host switching establishing during larval development that is irreversible/obligate after reaching the adulthood. Once again, the pre-acclimation to secondary hosts may indicate optimization of nutritional polyphenism, enabling rapid exploitation of new favorable habitats by the population. Altogether these observations suggest that the genetic programs needed to exploit the new host are probably switched on before the contact to the new host and even before take-off. The temporal sequence between wing development (first and second larval instar), pre-acclimation to secondary hosts (third larval instar) and preference switching (fourth larval instar) may indicate a finely regulated cascade of molecular mechanisms underlying nutritional polyphenism, which remains to be elucidated.

## 5. Conclusions

To clarify the critical periods of the nutritional polyphenism in *R. padi*, we studied two main steps of this process, the induction of the migrant morph and the establishment of diet change. We demonstrated that the cues from the plant maturity are key factors influencing the development of spring emigrants. We also identified that the window of perception occurs during the first and second larval stages enabling the development of emigrant within the same generation. In addition, the acclimation to the new host is gradually established along with the wing development. The current study thus documents, for the first time in host-alternating aphid species, the gradual and irreversible establishment of nutritional polyphenism. These findings highlight the remarkable capacity of *R. padi* to sense environmental cues, particularly those associated with plant-emitted signals from both primary and secondary hosts. The perception of plant cues constitutes a pivotal aspect of *R. padi’*s ecology, influencing its dispersal patterns, reproductive strategies, and population dynamics. From these findings, we aim to identify the plant compounds and the underlying molecular mechanisms that enable the host alternation in *R. padi*.

## Supporting information

Supplemental Statistical analysis

## Supplementary data

Supplementary informations are available online and include the following: Figure S1. **Figure S1**. The life cycle of *Rhopalosiphum padi*, a host-alternating species; **Figure S2**. Experimental set-up of the wing induction experiment; **Statistical analysis**. Detailed descriptions of the statistical tests, including models used, significance levels, and coefficients for each tested condition; **Table S1**. The GLMM results were utilized to assess the dependence of wheat preference on various stages of *R. padi*; **Table S2**. The Cox hazard model results were employed to examine whether the stage or morph transferred onto wheat influences the survival rate of *R. padi;* **Table S3**. The Cox hazard model results were employed to examine whether the plant age influences the survival rate of *R. padi*.

## Author Contributions

Conceptualization, J-C. Simon, Gaël Le-Trionnaire A. Etier and R. Ghosh.; methodology, A. Etier and R. Ghosh, T. Vericel, R. Cloteau, G. Denis, M. Le Floch; Data analysis, A. Etier, R. Ghosh and Y. Outreman; investigation, A. Etier, R. Ghosh.; writing—original draft preparation, A. Etier, R. Ghosh.; writing—review and editing, J-C. Simon, G. Le-Trionnaire, Y. Outreman, A. Etier, R. Ghosh; supervision, J-C. Simon.; project administration, J-C. Simon; funding acquisition, J-C. Simon. All authors have read and agreed to the published version of the manuscript.”

## Funding

This research was funded by ERC Alterevo, grant number 30001959.

## Acknowledgments

This work is dedicated to Professor A.F.G. Dixon, a pioneer in aphid ecology and biology and a source of inspiration to many aphid researchers.

## Conflicts of Interest

The authors declare no conflict of interest.

## Notes

### Competing Interest Statement

The authors have declared no competing interest.

