## Supplemental Figure S1 for "Insights into the triggering factors and determination mechanisms of host alternation in aphids"

### Supplementary data for insights into the triggering factors and determination mechanisms of host alternation in aphids


**Figure S1. The life cycle of *Rhopalosiphum padi*, a host-alternating species.** The life cycle is divided between the primary host (*Prunus padus*) depicted in dark green, and the secondary hosts (herbaceous plants) shown in yellow. Solid arrows indicate a single generation separating different morphs, while dotted arrows denote potential multiple generations of the same morph preceding the development of a novel morph. A focus on the difference between developmental stages of fundatrigeniae and adult emigrants during the spring migration includes four larval stages, with the first two stages being visually similar. The third and fourth larval stages differ, particularly in the absence or presence of wing buds for fundatrigeniae or emigrant respectively

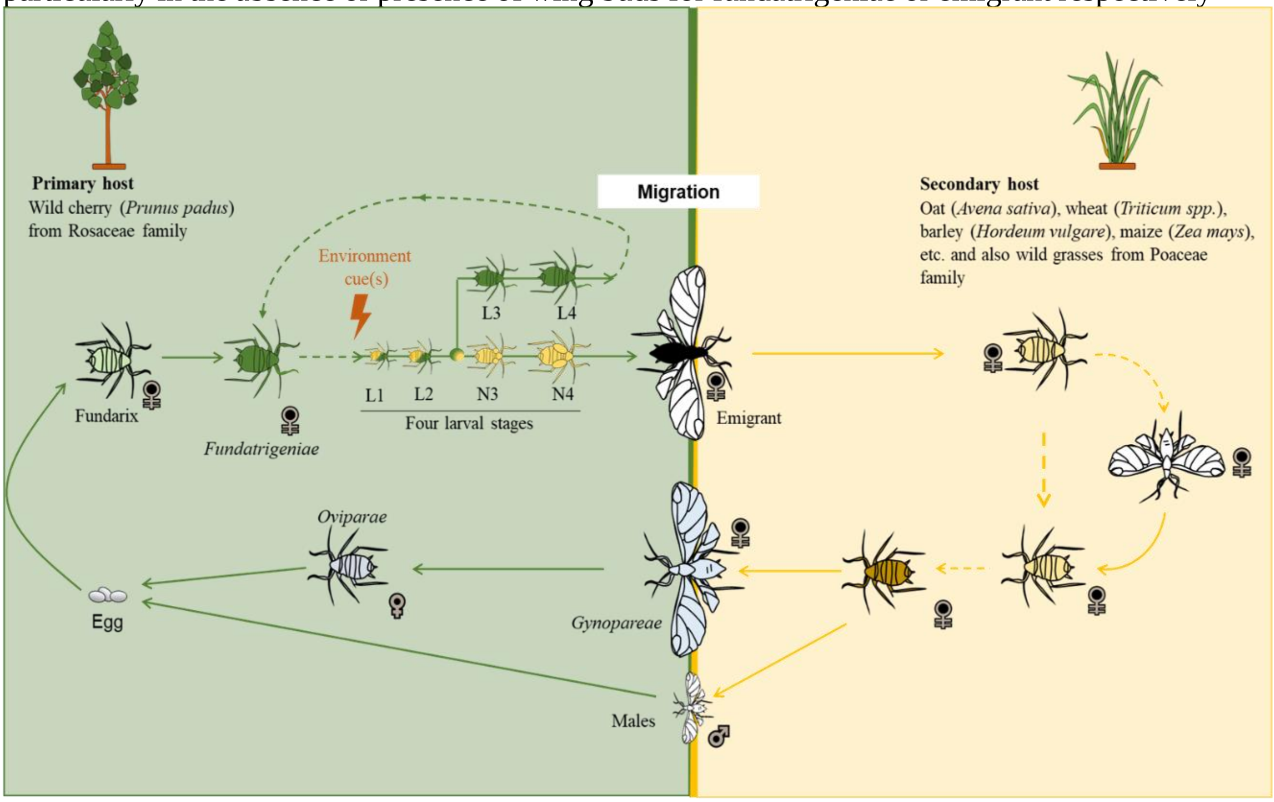
