## Supplemental Figure S2 for "Insights into the triggering factors and determination mechanisms of host alternation in aphids"

### Supplementary data for insights into the triggering factors and determination mechanisms of host alternation in aphids

**Figure S2. Experimental set-up of the wing induction experiment.** In red is represented the fundatrix kept on young leaves of *P. padus*. In light green, freshly born larvae from fundatrices referred here as Fundatrigeniae/Emigrant – first generation are either kept on young or mature leaves and their winged status were monitored after reaching the third larval stage. The wingless individuals from the first generation kept on young leaves were used to produced new larvae (dark green) that were deposited either on mature leaves after birth or after reaching the third larval instar. Here again the winged status was monitored after reaching the third larval instar.

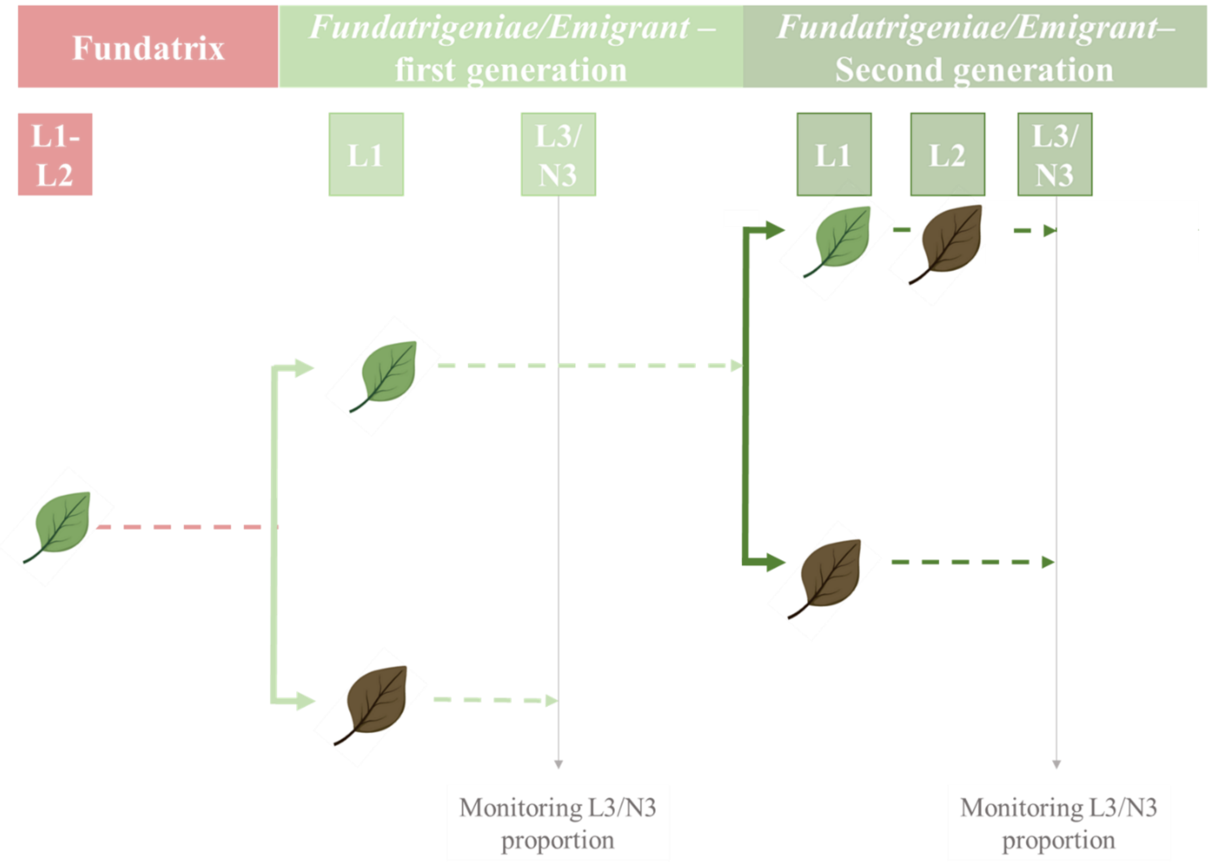
