## Supplemental Table S1 for "Insights into the triggering factors and determination mechanisms of host alternation in aphids"

### Supplementary data for insights into the triggering factors and determination mechanisms of host alternation in aphids

**Table S1. The GLMM results were utilized to assess the dependence of wheat preference on various stages of *R. padi*.** The first column of the table delineates the contrasts utilized to determine significant differences. The term preceding the 'vs' signifies the reference condition, while the term following it represents the condition being compared against the reference.

| Contrast | Estimate | SE | df | z.ratio | p.value |
|---|---|---|---|---|---|
| *Fundatrigeniae* vs L2 | 1.329 | 0.830 | Inf | 1.601 | 0.7497 |
| *Fundatrigeniae* vs N3 | -0.784 | 0.525 | Inf | -1.492 | 0.8122 |
| *Fundatrigeniae* vs N4 | -1.895 | 0.474 | Inf | -4.001 | 0.0016 |
| *Fundatrigeniae* vs Emigrant after the flight | -3.913 | 0.504 | Inf | -7.768 | <.0001 |
| *Fundatrigeniae* vs Emigrant before leaving *P. padus* leaves | -3.786 | 0.516 | Inf | -7.341 | <.0001 |
| *Fundatrigeniae* vs Wingless adult (wheat) | -7.237 | 1.090 | Inf | -6.638 | <.0001 |
| *Fundatrigeniae* vs Winged adult (wheat) | -6.582 | 0.830 | Inf | -7.934 | <.0001 |
| L2 vs N3 | -2.113 | 0.779 | Inf | -2.712 | 0.1187 |
| L2 vs N4 | -3.224 | 0.745 | Inf | -4.327 | 0.0004 |
| L2 vs Emigrant after the flight | -5.242 | 0.765 | Inf | -6.856 | <.0001 |
| L2 vs Emigrant before leaving *P. padus* leaves | -5.116 | 0.773 | Inf | -6.621 | <.0001 |
| L2 vs Wingless adult (wheat) | -8.567 | 1.233 | Inf | -6.950 | <.0001 |
| L2 vs Winged adult (wheat) | -7.912 | 1.010 | Inf | -7.837 | <.0001 |
| N3 vs N4 | -1.111 | 0.377 | Inf | -2.947 | 0.0636 |
| N3 vs Emigrant after the flight | -3.129 | 0.414 | Inf | -7.555 | <.0001 |
| N3 vs Emigrant before leaving *P. padus* leaves | -3.003 | 0.429 | Inf | -7.002 | <.0001 |
| N3 vs Wingless adult (wheat) | -6.454 | 1.052 | Inf | -6.135 | <.0001 |
| N3 vs Winged adult (wheat) | -5.799 | 0.779 | Inf | -7.448 | <.0001 |
| N4 vs Emigrant after the flight | -2.018 | 0.346 | Inf | -5.828 | <.0001 |
| N4 vs Emigrant before leaving *P. padus* leaves | -1.892 | 0.364 | Inf | -5.201 | <.0001 |
| N4 vs Wingless adult (wheat) | -5.343 | 1.027 | Inf | -5.202 | <.0001 |
| N4 vs Winged adult (wheat) | -4.688 | 0.745 | Inf | -6.295 | <.0001 |
| Emigrant after the flight vs Emigrant before leaving *P. padus* leaves | 0.126 | 0.402 | Inf | 0.314 | 1.0000 |
| Emigrant in flight vs Wingless adult (wheat) | -3.325 | 1.041 | Inf | -3.193 | 0.0306 |
| Emigrant after the flight vs Winged adult (wheat) | -2.670 | 0.764 | Inf | -3.493 | 0.0112 |

| | | | | | |
|---|---|---|---|---|---|
| Emigrant on *P. padus* vs Wingless adult (wheat) | -3.451 | 1.047 | Inf | -3.295 | 0.0220 |
| Emigrant on *P. padus* vs Winged adult (wheat) | -2.796 | 0.772 | Inf | -3.620 | 0.0071 |
| Wingless adult (wheat) vs Winged adult (wheat) | 0.655 | 1.232 | Inf | 0.531 | 0.9995 |
