## Supplemental Table S2 for "Insights into the triggering factors and determination mechanisms of host alternation in aphids"

### Supplementary data for insights into the triggering factors and determination mechanisms of host alternation in aphids

**Table S2. The Cox hazard model results were employed to examine whether the stage or morph transferred onto wheat influences the survival rate of *R. padi*.** The first column of the table denotes the contrasts utilized to assess significant effects. The term preceding the 'vs' indicates the reference condition, while the term following it represents the condition being compared against the reference**.**

| Contrast | coef | exp(coef) | se(coef) | z | p |
|---|---|---|---|---|---|
| L1 vs L2 | -0.2736 | 0.7607 | 0.2515 | -1.088 | 0.27663 |
| L1 vs N3 | -0.9396 | 0.3908 | 0.2664 | -3.527 | 0.00042 |
| L1 vs N4 | -1.5761 | 0.2068 | 0.3209 | -4.912 | 9.01e-07 |
| L1 vs Emigrant before leaving *P. padus* | -1.7605 | 0.1720 | 0.3394 | -5.188 | 2.13e-07 |
| L1 vs Emigrant after the flight | -2.2154 | 0.1091 | 0.3962 | -5.591 | 2.25e-08 |
| L2 vs L1 | 0.2736 | 1.3146 | 0.2515 | 1.088 | 0.2766 |
| L2 vs N3 | -0.6661 | 0.5137 | 0.2716 | -2.452 | 0.0142 |
| L2 vs N4 | -1.3025 | 0.2719 | 0.3247 | -4.011 | 6.03e-05 |
| L2 vs Emigrant before leaving *P. padus* | -1.4869 | 0.2261 | 0.3428 | -4.338 | 1.44e-05 |
| L2 vs Emigrant after the flight | -1.9419 | 0.1434 | 0.3991 | -4.866 | 1.14e-06 |
| N3 vs L1 | 0.9396 | 2.5590 | 0.2664 | 3.527 | 0.00042 |
| N3 vs L2 | 0.6661 | 1.9466 | 0.2716 | 2.452 | 0.01421 |
| N3 vs N4 | -0.6364 | 0.5292 | 0.3341 | -1.905 | 0.05679 |
| N3 vs Emigrant before leaving *P. padus* | -0.8209 | 0.4400 | 0.3516 | -2.335 | 0.01955 |
| N3 vs Emigrant after the flight | -1.2758 | 0.2792 | 0.4066 | -3.138 | 0.00170 |
| N4 vs L1 | 1.5761 | 4.8359 | 0.3209 | 4.912 | 9.01e-07 |
| N4 vs L2 | 1.3025 | 3.6785 | 0.3247 | 4.011 | 6.03e-05 |
| N4 vs N3 | 0.6364 | 1.8897 | 0.3341 | 1.905 | 0.0568 |
| N4 vs Emigrant before leaving *P. padus* | -0.1845 | 0.8316 | 0.3934 | -0.469 | 0.6392 |
| N4 vs Emigrant after the flight | -0.6394 | 0.5276 | 0.4432 | -1.443 | 0.1492 |
| Emigrant before leaving *P. padus* leaves vs L1 | 1.7605 | 5.8154 | 0.3394 | 5.188 | 2.13e-07 |
| Emigrant before leaving *P. padus* leaves vs L2 | 1.4869 | 4.4236 | 0.3428 | 4.338 | 1.44e-05 |
| Emigrant before leaving *P. padus* leaves vs N3 | 0.8209 | 2.2725 | 0.3516 | 2.335 | 0.0195 |
| Emigrant before leaving *P. padus* leaves vs N4 | 0.1845 | 1.2026 | 0.3934 | 0.469 | 0.6392 |

| | | | | | |
|---|---|---|---|---|---|
| Emigrant before leaving *P. padus* leaves vs Emigrant after the flight | -0.4549 | 0.6345 | 0.4564 | -0.997 | 0.3189 |
| Emigrant after the flight vs L1 | 2.2154 | 9.1654 | 0.3962 | 5.591 | 2.25e-08 |
| Emigrant after the flight vs L2 | 1.9419 | 6.9718 | 0.3991 | 4.866 | 1.14e-06 |
| Emigrant after the flight vs N3 | 1.2758 | 3.5816 | 0.4066 | 3.138 | 0.0017 |
| Emigrant after the flight vs N4 | 0.6394 | 1.8953 | 0.4432 | 1.443 | 0.1492 |
| Emigrant after the flight vs Emigrant before leaving *P. padus* | 0.4549 | 1.5761 | 0.4564 | 0.997 | 0.3189 |
