## Supplemental Table S3 for "Insights into the triggering factors and determination mechanisms of host alternation in aphids"

### Supplementary data for insights into the triggering factors and determination mechanisms of host alternation in aphids

**Table S3. The Cox hazard model results were employed to examine whether the plant age influences the survival rate of *R. padi*.** The first column of the table denotes the contrasts utilized to assess significant effects. The term preceding the 'vs' indicates the reference condition, while the term following it represents the condition being compared against the reference**.**

| **Contrast** | **coef** | ***Exp(coef)*** | **Se(coef)** | **z** | **p-val** |
|---|---|---|---|---|---|
| Wheat vs young leaves | 1.4362 | 0.4042 | 0.4042 | 3.553 | 0.000381 |
| Wheat vs mature leaves | 1.7976 | 0.3991 | 0.3991 | 4.504 | 6.66e-06 |
| Young leaves vs mature leaves | 0.3614 | 1.4354 | 0.2621 | 1.379 | 0.167853 |
| Young leaves vs wheat | -1.4362 | 0.2378 | 0.4042 | -3.553 | 0.000381 |
| Mature leaves vs young leaves | -0.3614 | 0.6967 | 0.2621 | -4.504 | 0.168 |
| Mature leaves vs wheat | -1.7976 | 0.1657 | 0.3991 | -1.379 | 6.66e-06 |
