## Supplemental Statistical analysis for "Insights into the triggering factors and determination mechanisms of host alternation in aphids"

### Supplementary data for insights into the triggering factors and determination mechanisms of host alternation in aphids

**Statistical analysis**.

To investigate the effect of generation exposure to the leaf maturity signal on winged induction, we employed a Generalized Linear Model (GLM). Specifically, we analyzed the presence or absence of the wings as a function of the leaf age. The explanatory variable in the model was the maturity of the leaves to which the generation of aphid was exposed. The response variable was the binary outcome of the wings' presence or absence, which was analysed using a binomial GLM with a logit link function. The logit link function was chosen to model the probability of wing induction as it is appropriate for binary outcome variables. Models were fitted using the "glm" function in R. The significance was assessed using a likelihood ratio test. This test was used to study a statistically significant effect of the leaf maturity on the wing induction depending on the generation. If a significant effect was observed, the model coefficients were examined to interpret the direction and magnitude of the effect of the leaf maturity on wing induction.

To analyze the effect of the stage L1 or L2 exposure to the leaf maturity in the emigrant's induction a GLM was performed. Explanatory variable was the age of the leaves. The presence or absence of the wings were analysed as a function of the leaf age. The explanatory variable in the model was the maturity of the leaves to which the stages L1 or L2 were exposed. The response variable was the binary outcome of the wings' presence or absence, which was analysed as previously detailed for the GLM. The likelihood ratio test was used to study a statistically significant effect of the leaf maturity on the wing induction depending on the stages. If a significant effect was observed, the model coefficients were examined to interpret the direction and magnitude of the effect of the leaf maturity on wing induction.

Generalized linear mixed models (GLMM) with a binomial distribution were used to test whether the different stages/morphs had an effect on the response variable (wheat choice). We used Petri dish as a random effect term. Pairwise comparisons were used to determine differences between the different stages/morphs and the number of individuals with choice and whose choice was wheat. When a significant effect of one of the main factors was detected or when an interaction between factors was significant, a pairwise comparison using least-squares means (package R: “emmeans”) (*p* value adjustment with Tukey method) was used to test for differences between treatments (Searle, Speed, & Milliken, 1980).

A Univariate Cox proportional hazards model was employed to examine the association between the developmental stages of emigrants and time to death after transfer onto wheat as secondary host. Analyses were conducted using the “survival” package in R. Kaplan Meier curves were used to visualize survival rate over the period of the assay and to estimate survival probabilities.

A Univariate Cox proportional hazards model was carried out to study the association between the survival of emigrants on either young or mature leaves or wheat to test the reversibility of the nutritional polyphenism in *R. padi*. Analyses and visualization were performed as previously described.
